# Dynamics of nonlinguistic statistical learning: From neural entrainment to the emergence of explicit knowledge

**DOI:** 10.1101/2020.07.30.228130

**Authors:** Julia Moser, Laura Batterink, Yiwen Li Hegner, Franziska Schleger, Christoph Braun, Ken A. Paller, Hubert Preissl

**Affiliations:** IDM/fMEG Center of the Helmholtz Center Munich at the University of Tübingen, University of Tübingen, German Center for Diabetes Research (DZD), Tübingen, Germany; Graduate Training Centre of Neuroscience, International Max Planck Research School, University of Tübingen, Tübingen, Germany; Western University, Department of Psychology, Brain and Mind Institute, London, ON, Canada; MEG Center, University of Tübingen, Tübingen, Germany; Center of Neurology, Department of Neurology and Epileptology, Hertie-Institute for Clinical Brain Research, University of Tübingen, Tübingen, Germany; CIMeC, Center for Mind/Brain Sciences, University of Trento, Trento, Italy; Northwestern University, Department of Psychology, Evanston, IL, USA; Department of Internal Medicine IV, University Hospital of Tübingen, Tübingen, Germany; Department of Pharmacy and Biochemistry, Interfaculty Centre for Pharmacogenomics and Pharma Research, University of Tübingen, Tübingen, Germany

**Keywords:** Statistical learning, MEG, auditory processing, implicit learning, explicit learning

## Abstract

Humans are highly attuned to patterns in the environment. This ability to detect environmental patterns, referred to as statistical learning, plays a key role in many diverse aspects of cognition. However, the spatiotemporal neural mechanisms underlying implicit statistical learning, and how these mechanisms may relate or give rise to explicit learning, remain poorly understood. In the present study, we investigated these different aspects of statistical learning by using an auditory nonlinguistic statistical learning paradigm combined with magnetoencephalography. Twenty-four healthy volunteers were exposed to structured and random tone sequences, and statistical learning was quantified by neural entrainment. Already early during exposure, participants showed strong entrainment to the embedded tone patterns. A significant increase in entrainment over exposure was detected only in the structured condition, reflecting the trajectory of learning. While source reconstruction revealed a wide range of brain areas involved in this process, entrainment in areas around the left pre-central gyrus as well as right temporo-frontal areas significantly predicted behavioral performance. Sensor level results confirmed this relationship between neural entrainment and subsequent explicit knowledge. These results give insights into the dynamic relation between neural entrainment and explicit learning of triplet structures, suggesting that these two aspects are systematically related yet dissociable. Neural entrainment reflects robust, implicit learning of underlying patterns, whereas the emergence of explicit knowledge, likely built on the implicit encoding of structure, varies across individuals and may depend on factors such as sufficient exposure time and attention.

## 1. Introduction

Living in a dynamically changing environment, humans and other animals are highly attuned to structure in their surroundings. They are able to extract relevant patterns from their surroundings using a process called *statistical learning* (Saffran et al., 1996a). Statistical learning is defined as the ability to extract the statistical properties of sensory input across time or space, and occurs automatically, incidentally and through mere passive exposure to the input (e.g. Frost et al., 2015; Schapiro and Turk-Browne, 2015; Siegelman et al., 2018a). The first experiment on statistical learning demonstrated that infants were able to extract embedded patterns from a continuous stream of speech by becoming sensitive to the transitional probabilities between neighboring syllables (Saffran et al., 1996a). This finding suggested that statistical learning plays an important role in language acquisition, and spurred a large body of additional work in this area, not only in developmental populations (e.g. Benitez et al., 2020; Fló et al., 2019; Pelucchi et al., 2009), but also in adults (e.g. Batterink and Paller, 2017; Cunillera et al., 2009; Saffran et al., 1996b). Importantly, many subsequent studies have shown that statistical learning is not restricted to language, but also occurs to nonlinguistic auditory (e.g. Gebhart et al., 2009; Saffran et al., 1999), visual (e.g. Bulf et al., 2011; Turk-Browne et al., 2005) and cross-modal stimuli (e.g. Cunillera et al., 2010; Paraskevopoulos et al., 2018).

One key benefit of using nonlinguistic stimuli to investigate statistical learning in the auditory domain is that it can be assumed that learners are “blank slates,” with learning not heavily influenced by prior knowledge. In a recent study, Siegelman and colleagues (2018b) demonstrated that statistical learning to linguistic stimuli (i.e., syllables) is strongly shaped by learners’ existing phonotactic knowledge and expectations about the input. To demonstrate this, they measured internal item consistency on one linguistic and two nonlinguistic (auditory and visual) statistical learning tasks. Results for items in the nonlinguistic tasks were highly correlated, while the linguistic task had a low internal item consistency (Siegelman et al., 2018b). Additionally, their experiments showed a strong within-subject correlation in performance in the two nonlinguistic tasks. This correlation across modalities was not present for a visual and a linguistic statistical learning task (Siegelman and Frost, 2015). These results suggest that basic, domain-general, statistical learning computations may be tested most reliably using stimuli that do not strongly invoke prior knowledge.

Statistical learning has many similarities to *implicit learning*, which is defined as “the capacity to learn without awareness of the products of learning” (Frensch and Rünger, 2003, p. 14). According to this view, implicit learning produces implicit knowledge – knowledge that can be expressed via a change in task performance, without requiring conscious retrieval. In contrast, explicit knowledge is accompanied by the awareness of memory retrieval, as assessed via recall and recognition tasks (Gabrieli, 1998; Schacter, 1987; Squire, 1987). Statistical learning can occur entirely implicitly. For example, even when learners fail to demonstrate explicit knowledge of the learned regularities, learning can be expressed on a performance-based task, as revealed by faster reaction times to more predictable elements (Batterink et al., 2015b). Similarly, using neuroimaging methods such as fMRI and MEG, neural evidence of learning has been observed even in the absence of above-chance recognition of the embedded regularities (Paraskevopoulos et al., 2012; Turk-Browne et al., 2009). Nonetheless, at least in adult learners, statistical learning is *usually* accompanied by explicit knowledge; that is, implicit and explicit knowledge may be accrued in parallel (Batterink et al., 2015a). In fact, the classic approach to studying statistical learning fundamentally relies on participants’ ability to explicitly recognize embedded regularities (e.g. Gebhart et al., 2009; Saffran et al., 1997; Saffran et al., 1996b). In this typical approach, participants are exposed to a stream of structured stimuli, and then complete a two alternative forced choice (2AFC) task, discriminating items from the stream and random items. Above-chance performance on this task is taken as evidence that statistical learning has occurred.

However, while behavioral discrimination measures provide a one-time snapshot of learners’ knowledge *after* the learning process, they neglect the temporal dynamics of learning—that is, the same level of performance on these tasks can originate from completely different learning trajectories (Siegelman et al., 2018a). In contrast, methods that involve monitoring *during* the learning period have been argued to provide a more complete picture of statistical learning. At the behavioral level, reaction time approaches can be used for such monitoring (Siegelman et al., 2018a). The use of neuroimaging methods to capture statistical learning is a possibly even more advantageous approach as such methods can track learning processes without the requirement of an overt behavioral task, and can also shed light on the neural mechanisms and neural structures involved in learning.

Both Electroencephalography (EEG) and Magnetoencephalography (MEG) have been used to track statistical learning during the learning period. A common neural marker in this context is the event-related N400 component, which shows an increase in amplitude during exposure to an artificial language compared to a random syllable stream (Cunillera et al., 2009; 2006). Another example is the mismatch negativity, which is elicited by stimuli with a low probability of occurrence, even when presented outside the focus of attention (e.g. Koelsch et al., 2016; Tsogli et al., 2019). Using MEG, Paraskevopoulos et al. (2012) demonstrated a mismatch response to part-triplets compared to triplets during exposure to a statistically regular stream, as early as 50ms after stimulus onset, even though participants’ post-exposure recognition was at chance level. This finding suggests that neural measures can be more sensitive indices of learning than post-exposure behavioral measures (see also Turk-Browne et al., 2009).

Another effective approach, which has a high signal-to-noise ratio and is especially well-suited to capturing the neural response to a continuous sensory stream, involves the measurement of *neural entrainment*. Neural entrainment refers to a property of the electromagnetic activity of the brain to resonate at the same frequency as an ongoing rhythmic stimulus. This neural response can be quantified by either investigating changes in the power (e.g. Buiatti et al., 2009; Farthouat et al., 2017) or inter-trial-phase coherence (ITC; e.g. Batterink and Paller, 2017) at the frequency of the stimulus and of larger embedded patterns in a statistical learning stream. Studies using this approach have demonstrated that neural entrainment to hidden patterns increases over the exposure period (Batterink and Paller, 2019; 2017) and also predicts performance on post-exposure behavioral tests (Batterink and Paller, 2019; 2017; Buiatti et al., 2009; Choi et al., 2020). Taken together, these results suggest that neural entrainment reflects the successful perceptual grouping of raw stimulus elements into cohesive units, which occurs as a consequence of statistical learning (e.g., syllables into words). Importantly, such methods allow for the quantification of statistical learning during the acquisition process, independent of participants’ later behavioral responses.

In addition to shedding light on the time course of learning (Batterink and Paller, 2019; 2017), neuroimaging studies have also revealed which areas in the brain are active during statistical learning tasks. Previous studies using functional magnetic resonance imaging (fMRI) and functional near infrared spectroscopy (fNIRS) have yielded mixed results. While most authors agree on the importance of the superior temporal cortex in statistical learning, other findings implicate the premotor cortex (Cunillera et al., 2009), the inferior frontal cortex (Abla and Okanoya, 2008; Karuza et al., 2013), or the supramarginal gyrus (McNealy et al., 2006). However, because fMRI and fNIRS rely on indirect measurement of neural activity (through blood oxygenation levels), these methods cannot directly capture the time-locking of neural activity to patterns in a sensory stream, as reflected by neural entrainment. To gain both temporal and spatial information, Farthouat et al. (2017) used MEG to assess nonlinguistic auditory statistical learning, presenting embedded tone triplets in a continuous stream. They detected an increase of power at the frequency of tone triplets from the third minute of exposure on, and were able to reconstruct this increase to the left posterior temporal sulcus and supplementary motor area. Nevertheless, participants’ behavioral responses in a 2AFC task were at chance level, precluding a direct link between neural responses and behavioral measures of learning (Farthouat et al., 2017).

These results, which include traditional recognition as well as neuroimaging measures, collectively highlight two important components of statistical learning: (1) the dynamic learning trajectory, which may occur even in the absence of explicit knowledge, and which can be revealed through sensitive neural measures and (2) explicit knowledge of the learned regularities, which seems to be present in some studies (or some participants) but not in others. Insight into how these two components are related is key to gaining a deeper understanding of the underlying dynamics and neural mechanisms of statistical learning. To this end, some previous studies have attempted to link neural responses with subsequent explicit knowledge. For example, in a linguistic statistical learning task, Karuza et al. (2013) showed that neural activation in the left inferior frontal gyrus was related to participants’ explicit ratings of words versus part-words. Similarly, Abla et al. (2008) reported that N400 amplitudes elicited by the first tone of a tone-triplet were highest within the first ∼7 min of exposure in participants classified as high learners based on 2AFC performance, compared to middle and low learners. In addition, as mentioned previously, several studies have found a positive association between neural entrainment during learning and subsequent performance on behavioral statistical learning tests (Batterink and Paller, 2019; 2017; Buiatti et al., 2009; Choi et al., 2020).

In the current study, we combined the strengths of multiple approaches in order to investigate statistical learning over time and space, and its reflection in behavior. We used a nonlinguistic task that is less likely to be influenced by learners’ prior knowledge. During exposure, we used MEG to monitor participants’ statistical learning to embedded triplet sequences, which provides sufficient temporal resolution to measure neural entrainment, and additionally allows us to examine the neural sources that are most relevant for this entrainment. Neural entrainment was quantified via the comparison of ITC at the triplet and tone frequencies between two stimulation conditions, in which tones were organized in repeating triplets, or in pseudorandomized order (Batterink and Paller, 2019; 2017). If statistical learning is present, we expect ITC at the triplet frequency to be higher in the structured condition compared to the random condition, reflecting stronger neural phase-locking to the embedded triplets in the structured condition. In addition, this phase-locking value at the triplet frequency is expected to increase over the course of the exposure period, reflecting the progression of learning (Batterink and Paller, 2019). Furthermore, we aimed to elucidate the neural sources of the neural entrainment, providing further insight into the core substrates of statistical learning. We expected the superior temporal cortex to be the major hub (Abla and Okanoya, 2008; Cunillera et al., 2009; McNealy et al., 2006).

After exposure, we tested participants’ learning of the tone triplets at the behavioral level by using a familiarity rating task, which is sensitive to explicit knowledge, and a speeded response time task, which assesses reaction times to individual tones presented in different positions within the triplets and captures implicit knowledge of the learned regularities (Batterink et al., 2015a). We expected triplets to be rated as most familiar and reaction times to tones that occurred in later triplet positions to be faster as a result of increased predictability (Batterink and Paller, 2017), providing behavioral evidence of statistical learning.

Lastly, we explored relations between performance in the behavioral tasks and neural entrainment. We incorporated results at both the sensor and source level across different phases of exposure to gain insight into the relationship between neural entrainment, the progression of learning, and explicit knowledge. We hypothesized that stronger neural entrainment to tone triplets should predict better behavioral performance on the post-learning tests.

## 2. Material and Methods

### 2.1. Participants

Participants were 24 healthy volunteers (12 male) between 20 and 37 years old (mean age 27.54 years, SD=9.96). All participants were right handed and had normal hearing abilities. Ten out of the 24 participants had no formal musical education (outside of lessons in school). Fourteen participants reported formal musical education, ranging from 1-16 years (M = 8; SD = 3.9). They received 10 € per hour for their participation. The local ethics committee of the Medical Faculty of the University of Tübingen approved the study (No. 231/2018BO1).

### 2.2. Materials and Design

Auditory stimulation consisted of 12 pure sinusoidal tones between 261.63 and 932.33Hz. The 12 tones corresponded to notes C, D, E, F#, G# and A# from the 4th and 5th octave of a standard piano (261.63Hz, 293.66Hz, 329.63Hz, 369.99Hz, 415.3Hz, 466.16Hz, 523.25Hz, 587.33Hz, 659.26Hz, 739.99Hz, 830.61Hz, 932.33Hz).The tones were organized into two different types of sequences, one so-called “structured” condition and one “random” condition (Figure 1). In both conditions, the 12 tones were grouped into triplets composed of three tones that never spanned more than one octave, allowing for better perceptibility. The structured condition consisted of only four triplets, which repeated over the course of exposure. In contrast, in the random condition, composition of the individual triplets changed constantly throughout the exposure block, resulting in a pseudo-random stream of tones. The random condition was composed of ever-changing triplets in order to better equate acoustic similarity in both the structured and random conditions. In both conditions, the triplets were presented in pseudorandom order, with the constraint that neither the same tone, nor the same triplet, could repeat consecutively. Within the structured condition, to control for a possible effect of the position of each individual tone within a triplet, three different structured sequences were created and presented counterbalanced across participants. Across the three counterbalanced sequences, tones within the triplets were the same but each tone occurred in a different triplet position (first, second or third; e.g. C-F#-E, E-F#-C and F#-C-E). In both conditions, each tone occurred an equal number of times. Each sequence consisted of 2400 tones. Tones had a duration of 300 ms and were presented every 333 ms (i.e., with an inter-tone interval of 33 ms), yielding a total duration of 13.32 minutes per stimulation block.

**Figure 1.**
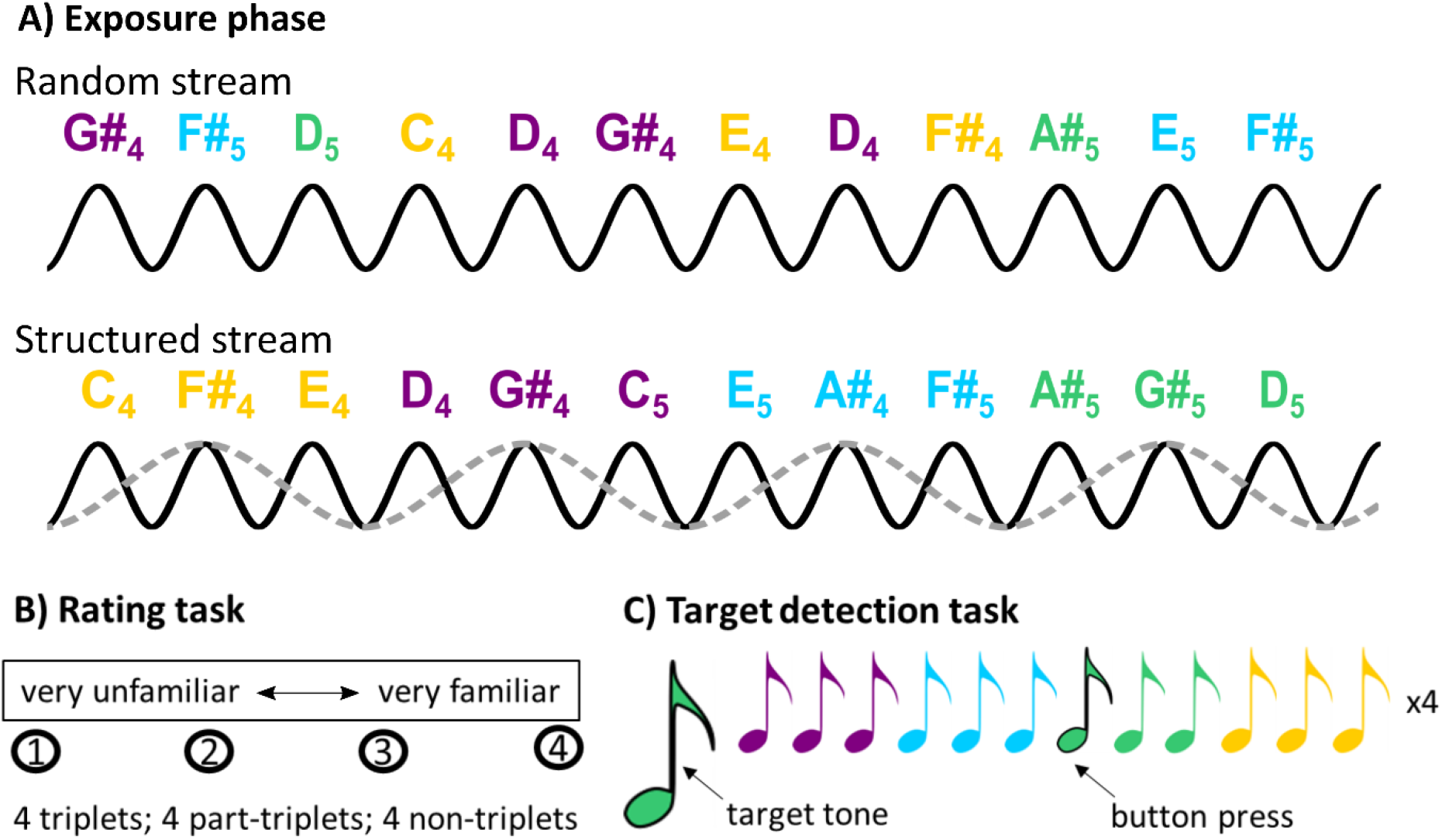
Summary of the experimental protocol. A) Auditory stimulation and hypothesized neural entrainment effects, as measured by MEG. The notes represent the different tones, the black sine waves show the frequency of tone presentation and the grey sine wave shows the frequency of triplets. If statistical learning of the underlying triplets occurs in the structured condition, stronger neural entrainment at the triplet frequency (grey dotted line) is expected relative to the random condition. B) Rating task performed after the exposure phase in the structured condition. Four triplets, four part-triplets and four non-triplets were rated on a scale from 1 (very unfamiliar) to 4 (very familiar). C) Target detection task performed after the rating task. Participants were presented with a target tone, which occurred four times within a stream of 36 triplets. Detection was indicated by a button press, and reaction time was measured.

Following the exposure phase, participants performed two behavioral tasks, designed to measure explicit and implicit aspects of statistical learning of the structured sequence (cf. Batterink and Paller, 2017). In the explicit rating task, participants were asked to rate 12 items for familiarity on a scale from 1 (very unfamiliar) to 4 (very familiar). Four of the items were triplets that were previously presented in the structured condition, four were part-triplets that consisted of a pair of tones from one of the triplets and a third tone that did not belong to the triplet, and four were non-triplets that were never presented in the structured block. In the implicit target detection task, participants were asked to detect target tones that occurred within short tone sequences that followed the same triplet structure as the original structured stimulation block. For each trial, participants were presented with a target tone, followed by a short stream of tones. Each stream consisted of four repetitions of the four tone triplets from the structured condition (presented with a slightly longer inter-tone interval of 66 ms to facilitate the task), yielding a total of four targets per stream. A total of 36 streams were presented, with each tone serving as the target three times (48 syllables per triplet position). Target detection was indicated by a button press, with both speed and accuracy emphasized in the instructions.

### 2.3. Procedure

Data were recorded using a 275-sensor, whole-head MEG system (VSM Medtech, Port Coquitlam, Canada) installed in a magnetically shielded room (Vakuumschmelze, Hanau, Germany). Auditory stimulation was produced by a loudspeaker outside of the shielded room and conducted through tubes into the shielded room. Tones were presented through earplugs connected to the tubes at an intensity of 70dB.

After arrival at the MEG laboratory, participants were fully informed about the experimental procedure and signed a consent form to confirm their voluntary participation. They filled out a short questionnaire, and completed a short hearing assessment with a screening audiometer (Hortmann Neuro-Otometrie Selector 20 K) to confirm normal hearing. Participants then changed into metal-free clothes. They were seated in a height-adjustable chair and instructed to fixate on a cross displayed in front of them during the whole recording. Participants were told that they would hear sounds through their earplugs, but that they had no particular task to perform related to those sounds. The MEG signal was recorded continuously, with a sampling rate of 585.94Hz.

During the exposure phase, participants listened to the random condition first, followed by the structured condition. We made the decision to consistently present the random condition first because we did not want participants to transfer or apply knowledge and expectations accrued during the structured block to the random block. The selected order thus avoids the possibility that participants may superimpose a triplet parsing scheme or otherwise explicitly search for patterns in the random condition. Two short breaks were included within each exposure block. After both exposure blocks, participants performed the rating task, followed by the target detection task. The target detection task was preceded by a short practice trial, with syllables rather than tones, to familiarize participants with the task. Both behavioral tasks were performed in the MEG chair to keep the environment consistent, but MEG was not recorded during these tasks. Finally, after both tasks were completed, participants were asked whether they heard any difference between the two exposure blocks.

### 2.4. Data Analysis

For analysis of the MEG data, MATLAB version 2016b (The MathWorks, Natick, MA) and the MATLAB-based open-source software package Fieldtrip (Oostenveld et al., 2011) were used.

### 2.4.1. Preprocessing

MEG data were band-pass filtered between 0.1-30Hz. Channels containing a high level of noise (overall root mean square (RMS) > 1 pT or no signal (overall RMS of 0.01 aT) were removed. For the main analysis of neural entrainment across the exposure period, data were time-locked to the onset of each triplet and extracted into nonoverlapping epochs containing 12 triplets (12000 ms)(cf. Batterink and Paller, 2017). This process led to a total of 64 epochs for evaluation (800 triplets formed by the 2400 stimuli, resulting in 67 epochs with 12 triplets, excluding 1 “partial” epoch near the end of each stimulation block to ensure an equal epoch length of 12000 ms). Epochs were corrected for a 32 ms sound output delay, and epochs containing artifacts with an amplitude over 4 pT were removed. This led to an average of 58.77 remaining epochs (SD=7.89). For the more fine-grained analysis of entrainment over time, data were extracted into epochs overlapping for 11/12 of their length in order to increase temporal resolution (cf. Batterink and Paller, 2017). This process led to a total of 766 epochs for evaluation (800 triplets formed by the 2400 stimuli, resulting in 799 overlapping epochs, excluding 11 epochs per block near the end of a stimulation block to ensure an equal epoch length of 12000 ms). After artifact rejection, an average of 703.19 epochs remained (SD=94.83).

### 2.4.2. Computation of inter-trial phase coherence (ITC)

We quantified overall neural entrainment across frequencies and across the exposure period by computing ITC across the nonoverlapping epochs. ITC is a measure of event-related phase-locking, with ITC values ranging from 1, reflecting completely phase-locked activity, to 0, indicating activity that is completely phase random with respect to the event of interest. Higher ITC values indicate more consistency in the phase of the signal across individual trials (in our case, epochs time-locked to triplet onsets). To the extent that statistical learning occurs, we expected to observe higher phase-locking at the triplet frequency, reflecting greater neural entrainment to the underlying triplet structure (Figure 1). ITC was computed using a fast Fourier transform with Hanning windows.

To analyze the temporal trajectory of neural entrainment in a fine-grained way, as a neural index of learning over time, we computed ITC values on “bundles” of 12 consecutive overlapping epochs (e.g., overlapping epochs 1-12, 13-24, 25-36, …755-766). As ITC values show greater fluctuation when computed over fewer trials, the resulting time course was smoothed over 5 bundles and the first and last bundle were omitted. This resulted in 61 bundles that represent the time course of ITC over the whole recording. For some participants, epochs were removed during artifact rejection reducing the total number of bundles. In this case, the participant’s remaining bundles were classified according to the closest temporal bundle position relative to the group average. Note that the use of overlapping epochs for this bundle analysis artificially increases the ITC values at the frequency of the overlap (in this case, at the triplet frequency), because the same data at this frequency occurs periodically and is considered repeatedly in calculations. Although this artifactual increase at the overlap frequency can complicate interpretation when computing ITC across frequencies, our time course analysis considers changes in ITC across time *within* a given frequency. Because this inflation is equally present across all bundles, the artifact does not affect the interpretation of ITC trajectories over time. We thus use overlapping epochs, rather than nonoverlapping epochs, for this analysis in order to increase the resolution of our temporal sampling, allowing for a more fine-grained analysis of the trajectory of learning over time.

### 2.4.3. Source Localization

To examine neural entrainment effects at the source level, we used the Dynamic Imaging of Coherent Sources (DICS) approach (Gross et al., 2001). In this approach, we created a surrogate signal, consisting of the combination of a 3Hz (tone frequency) and 1Hz (triplet frequency) sinusoidal and added it as an extra channel to the data. This allowed us to calculate coherence of the MEG signal with the tone and the triplet frequencies at the source level. A fast Fourier transform was calculated to obtain data in the frequency domain, which was then used as input for the DICS source analysis model. A common spatial filter for the structured and random conditions was calculated on the appended data to make conditions comparable. To project the source data, we used the ‘fsaverage’ brain, which is an average brain mesh provided by Freesurfer (Dale et al., 2012; Fischl et al., 1999), as a template brain and used SUMA (Saad and Reynolds, 2012) processing to obtain a decimated standard white/gray matter boundary mesh from which we could calculate a volume conduction head model with the ‘single shell’ method (Nolte, 2003). The MEG data were coregistered to the template head via the three fiducial points (nasion, left and right preauricular points). We obtained 2004 cortical surface vertices that were further used as MEG sources (cf. Li Hegner et al., 2018). For naming detected brain areas, we used the Destrieux et al. (2010) atlas.

### 2.4.4. Statistical Testing

Statistical tests were performed using R (R Core Team, 2019) and SPSS (IBM, 2017).

For the rating task, familiarity ratings were analyzed using a repeated-measures ANOVA with category (Triplet, Part-Triplet, Non-Triplet) as a within-participant factor. A linear contrast was used to test whether familiarity ratings decreased linearly across the three categories. Following previous studies (Batterink and Paller, 2019; 2017), a “rating score” was calculated for each participant, by subtracting the average score given for part-triplets and non-triplets from the average score given for triplets.

For the target detection task, for each participant, mean reaction times to detected targets were calculated within each triplet position (first, second, and third). Responses that did not occur within 0-1200 ms of a target were considered to be false alarms. The number of correctly detected tones and false alarms was quantified within each participant to get an estimate of task performance. To examine the hypothesis that reaction times should decrease linearly as a function of triplet position, reaction times were analyzed using a repeated-measures ANOVA with triplet position (initial, medial, final) as a within-participants factor, using a linear contrast. In an exploratory step, number of misses were compared in the same fashion.

At the sensor level, the interaction between condition (structured, random) and frequency (tone, triplet) was modeled with a mixed model in R (lme4; Bates et al., 2015), with ITC value as dependent variable and the two aforementioned predictors (condition, frequency). Participant was modeled as random intercept and the model was tested for significance with a type II ANOVA (lmerTest; Kuznetsova et al., 2017). Follow up analysis on ITC differences between conditions (structured versus random) at the triplet and tone frequencies were tested with a two-sided student t-test. Development of ITC over time was tested in an initial mixed effects model using ITC values within each bundle as the dependent variable. Predictors included condition (structured, random), sensor location (frontal, central, parietal, occipital, right temporal, left temporal), bundle number (as continuous predictor) and the full factorial interactions between these factors. Participant intercept was modelled as a random effect. Upon significant interactions with condition, post-hoc follow up analyses were conducted within each condition and factor of interest to characterize the time course of ITC over time. At the source level, statistical tests were performed in MATLAB with a cluster based permutation test (Maris and Oostenveld, 2007).

At the sensor level, relations between ITC and the rating score were explored with a stepwise multiple linear regression model, with the four key ITC variables (triplet frequency-structured, triplet frequency-random, tone frequency-structured, tone frequency-random) as predictors and rating score as the dependent measure. This analysis indicates which ITC variable(s) best predict the rating score, while removing variables that do not significantly contribute to the model. In the case of a significant model, we further explored the separate contribution of each ITC variable by calculating the Pearson’s correlation between each ITC variable and the rating score. At the source level, relations between triplet coherence and rating score were explored using the same type of model, with rating score and brain region (cluster) as predictors. Simple relations between triplet coherence in individual clusters and rating score were explored using Pearson correlations. To test whether the relation between the rating score and ITC differences between conditions strengthened or weakened over time, we computed the correlation values between the rating score and the ITC difference value (structured-random) at the triplet and tone frequency within each bundle. This produced an array of correlation values across time for each frequency (tone and triplet). A linear model with the correlation values as the dependent measure and time, frequency and their interaction as predictor variables was used to characterize the development of correlation values across time. As a follow-up, we also tested whether these values across time (within each frequency) showed a significant linear increase or decrease by using Pearson correlations.

Significance values were set to p<0.05. To account for the risk of false positives due to multiple comparisons, the source level analysis used a cluster-based correction with a cluster threshold at α=0.025, as the test was two-sided. For the bundle-by-bundle analysis of differences between conditions and the follow-up analyses regarding the relations between neural entrainment and behavior, significance values were adjusted with the Benjamini-Hochberg false discovery rate (FDR) procedure.

## 3. Results

### 3.1. Behavioral Results

In a subjective report after the experiment, 7 out of 24 participants reported that they did not hear any difference between the two conditions.

In the rating task, participants rated triplets (M = 3.07, standard error (SE) = 0.13) as more familiar compared to part-triplets (M = 2.91, SE = 0.12) and scrambled non-triplets (M = 2.79, SE = 0.12), as reflected by a significant linear effect (F(1,23) = 4.64, p = 0.042, *η*_*p*_^*2*^ = 0.17). The test for a category effect was only marginal (F(2,46) = 2.79, p = 0.075, *η*_*p*_^*2*^ = 0.11), showing that the effect of category on familiarity is small in magnitude for non-linguistic stimuli. The rating score was significantly above zero, providing evidence of above-chance learning (M = 0.23, SE = 0.10, 95% CI = 0.0089 – 0.44, t(23) = 2.15, p = 0.042).

In the target detection task, no significant effect of triplet position on reaction times was found (Triplet Position Effect: F(2,46) = 0.78, p = 0.44 *η*_*p*_^*2*^ = 0.033; linear contrast: F(1,23) = 0.64, p = 0.43, *η*_*p*_^*2*^ = 0.027). There was a significant effect of triplet position on accuracy, such that the number of misses (out of 48) increased with later triplet positions (Position Effect: F(2,46) = 16.2, p < 0.001, *η*_*p*_^*2*^ = 0.041; linear effect: F(1,23) = 36.7, p < 0.001, *η*_*p*_^*2*^ = 0.062). However, this effect was unexpected and in the opposite-to-predicted direction. Overall, participants performed poorly on the task, detecting only 63% of tones, compared to the syllable version of the task where 83-89% accuracy has been reported (Batterink & Paller, 2017, 2019). In addition, participants made an average of 102.5 false alarms (std = 49.4), which is much higher than in the linguistic version (∼12-19 false alarms, using the same total number of targets). Based on these results, we conclude that this version of target detection task, as implemented with tones rather than syllables, was too difficult to reveal significant learning effects. In the subsequent correlation analyses with MEG neural entrainment effects, only the rating score was used as a behavioral measure of learning.

### 3.2. Neural Entrainment at the Sensor Level

Across the exposure period and averaged across all MEG channels, ITC values showed a highly significant interaction between frequency (tone versus triplet) and condition (structured versus random; F(1,69) = 7.05, p = 0.010, ŋ _p_^2^ = 0.09). Critically, follow-up analyses showed that ITC at the triplet frequency was significantly higher in the structured compared to random condition (t(23) = 3.61, p_FDR_ = 0.003, d=0.74; Figure 2A). ITC at the tone frequency did not differ significantly between conditions (t(23) = -1.68, p_FDR_ = 0.106, d=-0.34).

**Figure 2.**
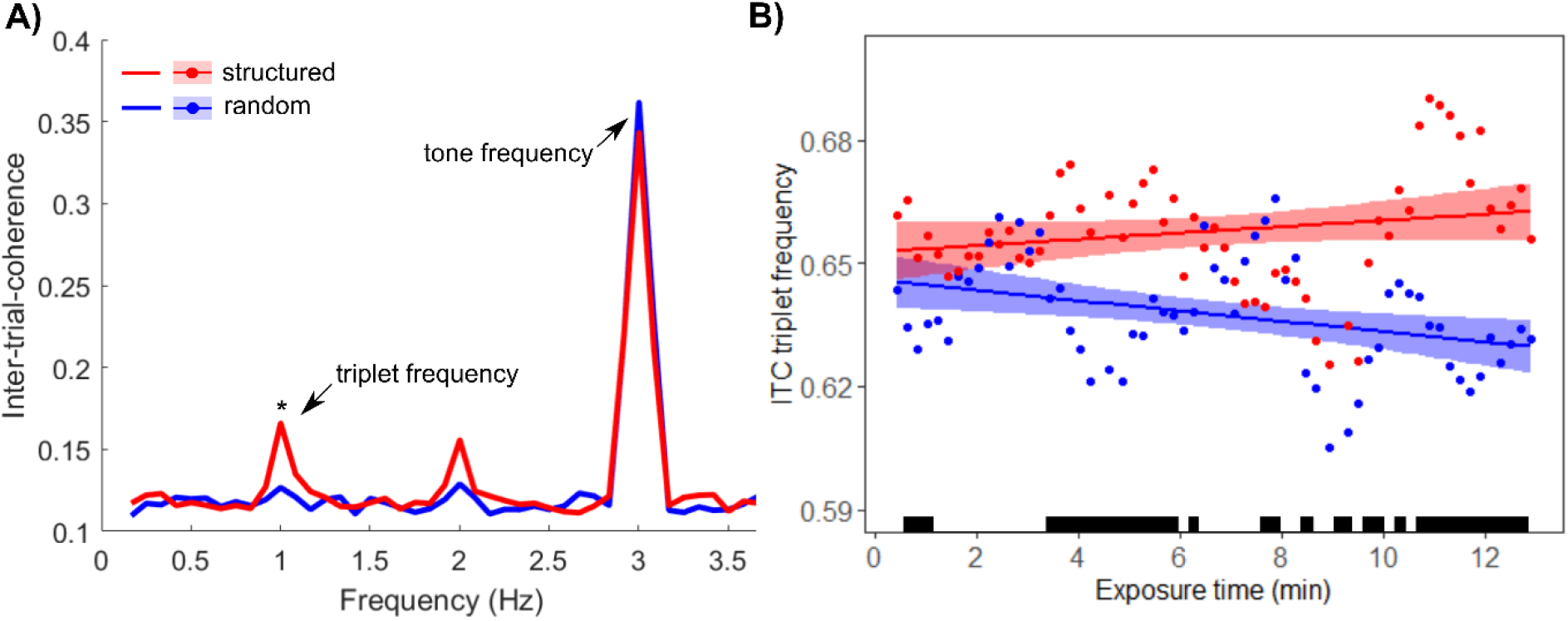
ITC differences between structured and random condition A) Inter-trial-coherence (ITC) at the whole brain level over the whole recording. Triplet frequency: 1Hz, Tone frequency: 3Hz. A significant difference was found between the structured and random conditions at the triplet frequency (* p<0.05). No difference was found at the tone frequency. B) Time course of ITC at the triplet frequency in the structured and random conditions, calculated in bundles over exposure time in minutes. Shaded areas represents 95% confidence interval of linear model; Black bars indicate time points at which the conditions significantly differed (34 out of 61 time points; p<0.05, FDR corrected).

A fine-grained, bundle-by-bundle analysis of ITC at the triplet frequency indicated that the time course of ITC differed significantly as a function of condition, with ITC in the structured condition showing a greater increase over time compared to the random condition (Condition x Time: F(1, 2657) = 14.5, p < 0.001, fixed effect estimate of interaction term = 0. 000428, SE = 0. 000112). Within the structured condition, a significant overall increase in triplet-ITC was revealed as the exposure period progressed (F(1,1332) = 4.69, p = 0.031; parameter estimate = 0.000164, SE = .000076). In contrast, within the random condition, a significant overall decrease in triplet-ITC was found as the exposure period progressed (F(1,1303) = 13.6, p < 0.001, parameter estimate = -0.000287, SE = 0.000078; Figure 2B).

Next, we investigated whether these effects varied as a function of sensor regions. The three-way interaction between time, region, and condition was not significant (F(5,15896) = 1.12, p = 0.35), suggesting that the relative increase over time in the structured condition compared to the random condition was similar across sensors. However, ITC showed significant differences in condition effects as a function of sensor region (Condition x Region: F(5,15896) = 2.25, p = 0.047), suggesting that overall condition effects were larger in some regions than others.

A follow-up Time x Region analysis of the structured condition alone replicated the overall effect of time across all sensors (F(1, 8001) = 15.5, p = 0.001, parameter estimate = 0.000166, SE = 0.000042). This increase over time did not differ significantly across the sensor regions (Time x Region: F(5,8001) = 1.58, p = 0.16). Similarly, in the random condition, a significant overall decrease in triplet-ITC was found as the exposure period progressed (F(1,7874) = 33.9, p < 0.001, parameter estimate = -0.000260, SE = 0.000045), which again did not differ across sensor regions (F(7872) = 1.57, p = 0.17). In addition, when compared directly to the random condition, ITC in the structured condition showed greater increases over time in each individual sensor region (all p values < 0.001). In sum, the increase in ITC over time is specific to the structured condition, suggesting this increase reflects statistical learning, rather than any nonspecific effects of stimulation over time.

To follow up on the significant Condition x Time interaction effect at the triplet frequency (reported above, p < 0.001), we further explored *when* during exposure a significant condition effect emerged. The goal of this analysis was to provide insight into the amount of exposure necessary for statistical learning to occur. For each of the 61 bundles, we tested a mixed effects model that included condition (structured, random) as a predictor, with participant intercept modelled as a random effect. As shown in Figure 2B, a total of 34 out of 61 bundles showed significantly greater ITC values in the structured condition relative to the random condition (p_FDR_ < 0.05). Interestingly, significant condition differences were already apparent in the earliest bundles (bundles 2-4), corresponding to the first ∼1 min of exposure. This finding indicates that neural entrainment to the embedded triplets emerges very rapidly, possibly after just a few exposures to the regularities.

### 3.3. Neural Entrainment at the Source Level

Clusters showing significant differences between the structured and random conditions in coherence with the surrogate signal at the triplet frequency (“triplet coherence”) are shown in Figure 3. The largest cluster spans over right temporo-frontal areas, including part of the superior temporal gyrus, the supramarginal and subcentral gyrus as well as part of the inferior frontal gyrus (cluster 1). On the left side, this homologous cluster is smaller, consisting of the superior temporal gyrus and part of the supramarginal and subcentral gyrus (cluster 3). Additionally, we found significant coherence differences between conditions in the left precentral gyrus together with parts of the middle and superior frontal gyri, spanning towards the back to parts of the postcentral gyrus and superior parietal lobe (cluster 3) and in the right superior parietal lobe (cluster 4). Structured and random conditions did not show any significant differences in coherence with the surrogate signal at the tone frequency (“tone coherence”).

**Figure 3.**
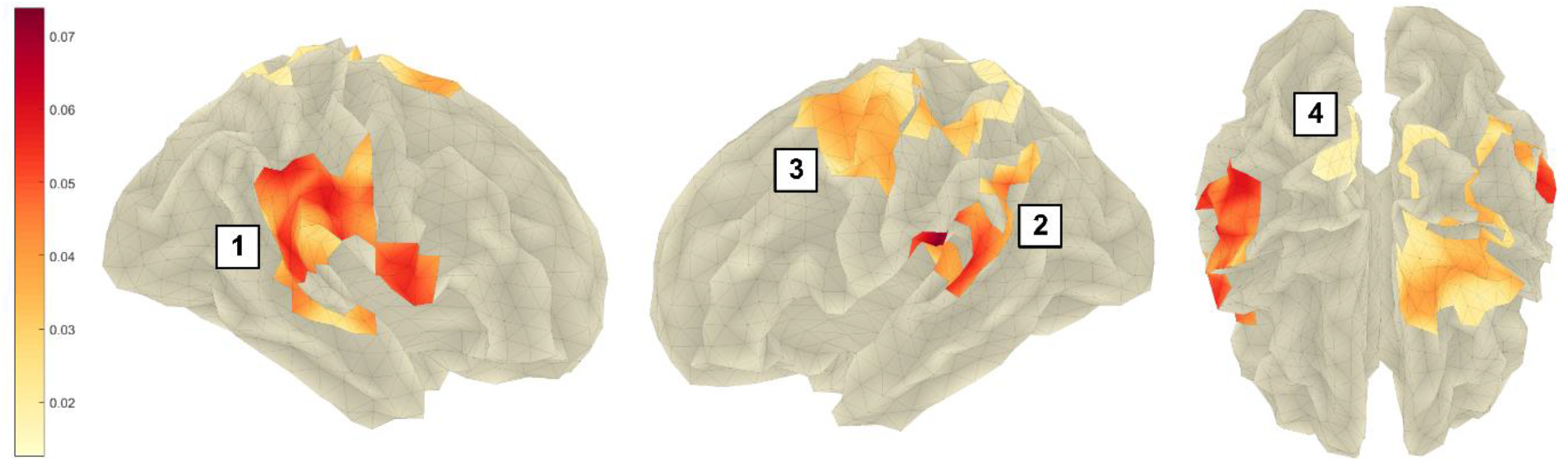
Areas where triplet coherence significantly (cluster threshold α<0.01) differed between structured and random conditions. Color gradient represents difference between triplet coherence in structured-random condition. For naming detected brain areas, we used the Destrieux et al. (2010) atlas. 1) right hemisphere: part of the superior temporal gyrus, supramarginal and subcentral gyrus, part of the inferior frontal gyrus; 2) left hemisphere: parts of the superior temporal, supramarginal and subcentral gyrus; 3) left hemisphere: precentral gyrus with parts of middle and superior frontal gyri, towards the back part of the postcentral gyrus and superior parietal lobe; 4) right superior parietal lobe.

### 3.4. Relationship Between Neural Entrainment and Behavior

#### 3.4.1. Neural Entrainment at the Sensor Level

A stepwise regression model indicated that when all 4 key ITC variables were entered in the model, only triplet-ITC in the structured condition significantly predicted the rating score (F(1,22) = 11.25, *p* = 0.003, r=0.58; Figure 4A). When each of the 4 variables was considered in isolation, tone-ITC in the structured condition as well as tone-ITC and triplet-ITC in the random condition showed a positive but non-significant relation with the rating score (r(22) = 0.38, p_FDR_ = 0.131; r(22) = 0.33, p_FDR_ = 0.160; r(22)=0.12, p_FDR_ = 0.573).

**Figure 4.**
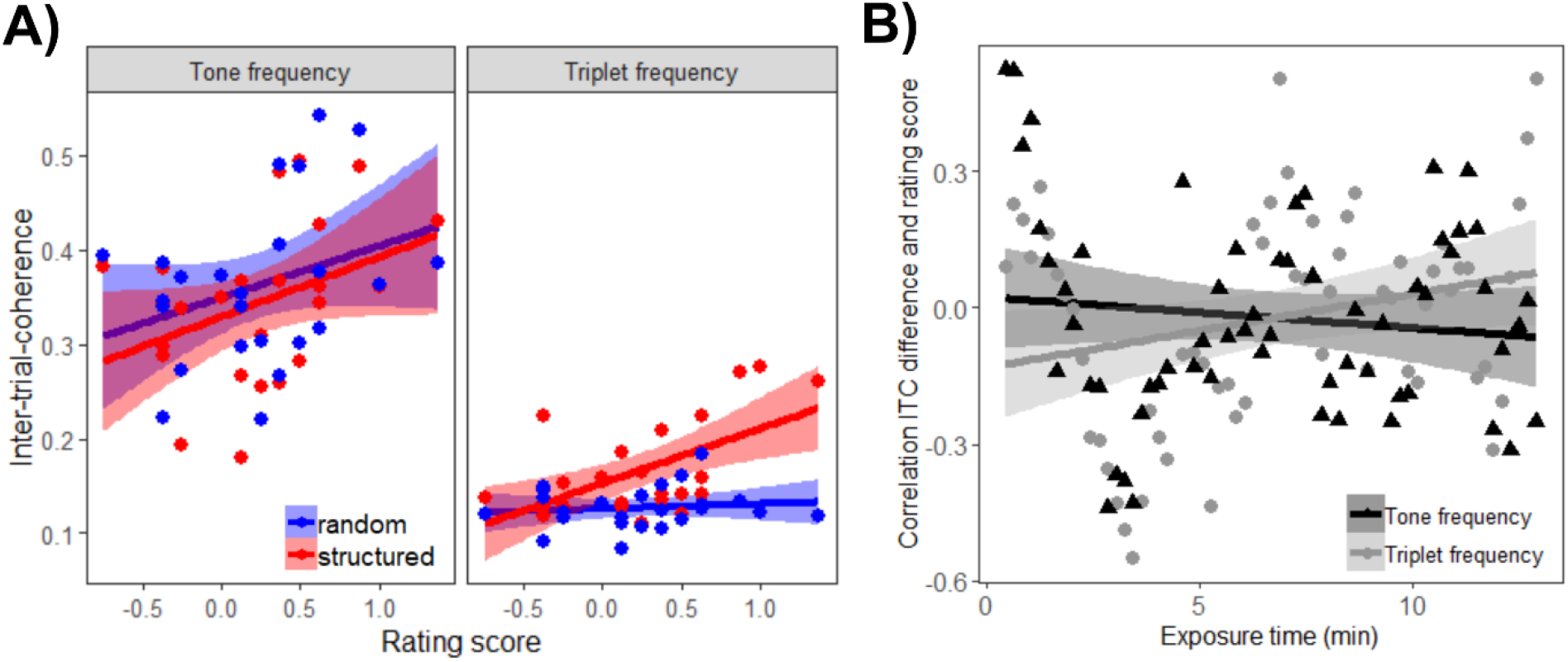
Relation between ITC and rating score. A) Relation between rating score and ITC at the tone frequency (left) and triplet frequency (right) in the structured and random conditions. Triplet ITC in the structured condition significantly correlated with rating score. B) Correlation between ITC difference value (structured-random) and rating score within each bundle over exposure time for triplet and tone frequency. Values showed a significant time x frequency interaction (p<0.05); Shaded areas represents 95% confidence interval of linear model.

To investigate how this relationship develops over time, for each bundle, we computed the correlation between rating score and the ITC difference between the structured and random condition at the (1) triplet frequency and (2) tone frequency. The development of these correlations over time significantly interacted with the frequency (F(1,118) = 4.41, p = 0.038, ŋ _p_^2^ = 0.04). At the triplet frequency, the correlation between rating score and ITC difference scores (structured – random) showed a tendency of increasing in strength (r(59) = 0.25, p_FDR_ = 0.101), whereas at the tone frequency this correlation seemed to decease in strength (r(59) = -0.12, p_FDR_ = 0.347; Figure 4B). However, following the significant interaction neither of these correlations did reached significance.

#### 3.4.2. Neural Entrainment at the Source Level

At the source level, the difference in triplet coherence between the structured and random conditions across all significant clusters (see Section 3.3) strongly correlated with the rating score (r(22) = 0.56, p = 0.004). A stepwise regression with all individual clusters entered as predictors revealed that cluster 3 (left precentral and parietal regions) mainly contributed to this correlation (F(1,22) = 2.14, p = 0.002, r = 0.60). Follow-up tests of individual clusters revealed another significant correlation in the right temporo-frontal cluster (r(22) = 0.53, p_FDR_ = 0.014; cluster 1), while the left temporal cluster showed a marginally significant relationship with the rating score (r(22) = 0.39, p_FDR_ = 0.079; cluster 2) and the right superior parietal cluster had no predictive value (r(22) = 0.24, p_FDR_ = 0.260; cluster 4).

## 4. Discussion

The present study showed that participants were able to learn nonlinguistic statistical regularities simply through exposure to the stimulus stream, in the absence of explicit instructions to detect the patterns. Participants demonstrated learning at both the behavioral and neural levels. At the behavioral level, participants rated triplets from the stimulus stream as significantly more familiar than non-triplets, revealing explicit knowledge of the learned regularities. At the neural level, learning was evidenced by robust neural entrainment to the embedded triplets across all sensors. This triplet entrainment was significantly greater in the structured condition compared to the random condition and emerged early on during exposure. In addition, triplet entrainment significantly increased over the course of exposure, reflecting the online trajectory of learning. Concerning the neural sources of these entrainment effects, a broad range of regions spanning temporal, frontal, and parietal cortices showed sensitivity to the hidden structure, which is generally in line with previous literature. Neural entrainment in these identified clusters also robustly predicted subsequent behavioral performance on the rating task. These results suggest that successful neural entrainment forms a basis for subsequent expression of explicit forms of knowledge.

### 4.1. Behavioral Results

As a group, participants showed significant behavioral evidence of learning as measured on the rating task, even though 7 out of 24 participants claimed that they did not observe any difference between random and structured blocks. Participants’ ratings distinguished among triplets, part-triplets, and non-triplets, although the differences in ratings were small and variable across participants. Linguistic statistical learning studies using a similar rating task have shown stronger learning effects (e.g. Batterink and Paller, 2017), which may be related to facilitation of learning enabled by verbal encoding strategies (Siegelman et al., 2018b). In contrast, nonlinguistic auditory statistical learning studies have reported both stronger (Abla et al., 2008; Abla and Okanoya, 2008; Gebhart et al., 2009; Saffran et al., 1999) and weaker, chance-level behavioral learning effects (Farthouat et al., 2017; Paraskevopoulos et al., 2012). These previous studies used a 2AFC task to test participants’ learning abilities, testing the contrast between triplets and either non-triplets (Abla et al., 2008; Abla and Okanoya, 2008) or part-triplets (Farthouat et al., 2017; Gebhart et al., 2009; Paraskevopoulos et al., 2012). Additionally, exposure time to the structured stream varied from 10 min (Farthouat et al., 2017) to 40 min (Gebhart et al., 2009). These variable results across studies emphasize that task difficulty (differentiating triplets from part-triplets or non-triplets) as well as exposure time likely play a crucial role in the recognition of tone structures. Therefore, even if a neural signature is visible, the ability to explicitly express knowledge acquired during statistical learning is not present automatically.

Effects in the target detection task were counterintuitive, showing a lower number of misses for the first tone in a triplet. Based on previous studies using the same task with language stimuli (e.g. Batterink et al., 2015b; Batterink et al., 2015a; Batterink and Paller, 2019, 2017) and abstract shapes (Kim et al., 2009; Turk-Browne et al., 2005), we had expected a facilitation for the third tone of a triplet, as it is most predictable. However, the low task accuracy, as well as subjective reports from participants, suggest that the task was too difficult to reveal expected learning effects. Moreover, memory for the specific tones may have been too weak to drive expectations for the third stimulus and support faster responding, as occurred in prior statistical learning studies with more complex stimuli. We speculate that the higher accuracy for the first tone of a triplet could have been caused by improved memory or improved perceptual separation for these less predictable tones, which could have in turn facilitated recognition. As the positions of individual tones in the structured condition were counterbalanced across participants, better accuracy for first tones cannot be due to stimulus-specific properties of certain tones.

### 4.2. Neural Entrainment

Neural entrainment to triplets was significantly enhanced in the structured compared to random tone streams. This condition difference was visible at the whole brain level with a relatively large effect size (d=0.74).

In the structured stream, ITC at the triplet frequency increased over the exposure period. In contrast, ITC in the random condition decreased over time, indicating that the observed increase in the structured condition is likely indicative of statistical learning, and cannot be attributed to nonspecific effects of auditory stimulation over time. The observed increase fits with previous neural entrainment results on linguistic statistical learning, where an increasing trajectory was reported over centro-frontal midline EEG electrodes (Batterink and Paller, 2019, 2017). Across studies, this increase in neural entrainment to the underlying triplets may reflect a shift in perception and encoding from individual stimulus units to more integrated items, which occurs gradually over the course of exposure. We also found that significant differences in triplet-ITC between structured and random conditions occurred very early during exposure (e.g., within the first 2-4 bundle included in the analysis, corresponding to approximately the first 35-70 seconds of exposure). This finding suggests that neural entrainment effects may emerge after just a few exposures to a given regularity. Similarly, a previous study by Barascud et al. (2016) who used repeating tone sequences of varying length (5-20 tones) demonstrated that participants were able to detect regularities after approximately one and a half repetitions of a given sequence. These effects may at least partially reflect a short-lived memory trace in auditory sensory memory, causing a change in the neural response to a recently encountered stimulus, similar to mechanisms that underlie the mismatch negativity (Näätänen et al., 2007).

An advantage over previous EEG-based neural entrainment studies of statistical learning (e.g., Batterink and Paller, 2017, 2019; Buiatti et al., 2009) is our use of MEG, which allowed us to localize specific cortical regions involved in learning. Overall, results are in line with previous statistical learning studies, while also revealing additional brain areas that have not been previously discussed in this context. Our localization analyses revealed a network of regions spanning the superior temporal, supramarginal, subcentral and inferior frontal cortex that showed robust differences in triplet coherence between the two conditions (Figure 3; clusters 1 and 2). These regions fit well with previous literature (Abla and Okanoya, 2008; Cunillera et al., 2009; Farthouat et al., 2017; McNealy et al., 2006). In both hemispheres, the clusters that span the superior temporal towards the inferior frontal cortex reflect the auditory processing hierarchy, and include Heschl’s gyrus, which is known to be important in pitch and melody perception (e.g. Patterson et al., 2002; Schneider et al., 2005). This most likely reflects a domain-specific component of auditory statistical learning, and suggests that even regions involved in early sensory processing may be sensitive to statistical regularities. The importance of the auditory cortex and inferior frontal gyrus for the early detection of auditory regularities has also been previously highlighted by Auksztulewicz et al. (2017), who used a dynamic causal model to explain the mechanism behind the early detection of repetitions described by Barascud et al. (2016).

The cluster spanning the precentral gyrus towards the superior frontal gyrus (cluster 3) is in line with findings by Cunillera et al. (2009) and Farthouat et al. (2017), who found the premotor cortex to be important for statistical learning. Based on these results, the authors argued for an audio-motor interface theory of speech learning, in which audio input is linked with motor representations of speech, thereby facilitating novel word learning (Cunillera et al., 2009). Converging evidence for this idea comes from a recent study by Assaneo and colleagues (2019), in which it was demonstrated that about half of participants spontaneously align their own speech output with a rhythmic syllable sequence. Remarkably, participants classified as high synchronizers not only showed higher brain-to-stimulus MEG synchrony over frontal areas, but also a better performance in a statistical language learning task compared to their low-synchronizing peers. Our results, as well as those of Farthouat and colleagues (2017), suggest that this audio-motor framework may also be applied to learning of nonlinguistic auditory sequences. In addition to memory for words, memory for tone patterns could also be mediated by the phonological working memory loop (Baddeley et al., 1998). Interestingly, the left lateralization of this precentral cluster, which Cunillera et al. (2009) connected to speech production, also holds for our nonlinguistic study as well as that of Farthouat and colleagues, suggesting that this left-lateralization is not strictly specific to linguistic processing.

Interestingly, the role of left fronto-motor areas has also been discussed in the context of prediction of upcoming speech. Using causal connectivity analysis, Park et al. (2020; 2015) demonstrated that delta range (1-4Hz) oscillations from left motor and frontal areas have a top-down modulatory influence on auditory cortices during natural speech processing. A further analysis examining coupling between brain beta frequency activity and speech delta phase showed a progression of speech prediction from higher order frontal areas to the auditory cortices (Park et al., 2020). Although we did not examine causal connectivity, similar predictive mechanisms may also be involved in the current study. Given the alignment in neural regions across Parks et al.’s studies and the current study, stronger neural entrainment in the left frontal motor regions may reflect prediction of upcoming predictable tones and could have a top-down impact on processing in auditory regions.

The clusters that we detected in the superior parietal cortex (clusters 4 and part of cluster 3) have not been reported in prior statistical learning literature. The superior parietal gyrus has been implicated in attention shifting (Vandenberghe et al., 2001). As clusters reflect differences in triplet coherence between structured and random conditions, a possible explanation for the involvement of the superior parietal gyrus could be an increase in attention to the triplets, once participants detected the regularity in the structured condition. The lack of regularity in the random condition could lead to decreased attention.

We observed bilateral clusters along the auditory processing stream, whereas some previous statistical learning studies found left hemispheric lateralization in these regions (Abla and Okanoya, 2008; Karuza et al., 2013; McNealy et al., 2006). However, both McNealy et al. (2006) and Karuza et al. (2013) used linguistic stimuli, which may be expected to result in greater left-lateralization compared to our nonlinguistic paradigm. Regarding lateralization for non-linguistic stimuli, previous results are inconsistent. Abla and Okanoya (2008) used a similar tone paradigm in an fNIRS study and reported a left-lateralization. In contrast, Farthouat et al. (2017) did not find a clear lateralization of statistical learning effects using MEG, and Janacsek et al. (2018) even emphasized the role of the right hemisphere, applying transcranial direct current stimulation during statistical learning. In sum, results of our study together with previous literature do not support the assumption of a lateralization of statistical learning per se. It is more likely that lateralization is a result of earlier sensory processing, influenced by acoustic features (e.g., high temporal resolution of speech and high spectral resolution of musical tones) as described by Zatorre and colleagues (2002).

### 4.3. Relationship Between Neural Entrainment and Behavior

Interestingly and as predicted, participants’ neural entrainment to the triplets in the structured condition was related to their behavioral learning outcomes, as measured by performance on the rating task. In the language domain, Ding et al. (2016) previously showed that neural tracking of phrases and sentences in speech depends on language comprehension, appearing only in participants who have knowledge of a given language (e.g., Mandarin). This finding demonstrates that neural entrainment is closely connected to high-level abstract knowledge and behavior. Further, as mentioned in the Introduction, a number of linguistic statistical learning studies have also a positive association between neural entrainment during learning and subsequent performance on behavioral tests of knowledge (Batterink and Paller, 2019; 2017; Buiatti et al., 2009; Choi et al., 2020). Along the same lines, our findings suggests that neural entrainment for tone-triplets is at least partially related to the emergence of explicit knowledge, with successful neural entrainment predicting the subsequent ability to explicitly recognize tone triplets. This brain-behavior correlation was observed at both the sensor and source level – that is, for triplet-ITC across all electrodes and for triplet coherence in the brain areas depicted in Figure 3. At the source level, this relationship was most pronounced in precentral/parietal areas in the left hemisphere (cluster3) followed by temporo-frontal areas in the right hemisphere (cluster1). The correlation with entrainment in temporal regions implicates domain-specific mechanisms, and suggests that areas involved in auditory processing also play a role in the formation of consciously available representations. The correlation with precentral regions points towards a possible role of the previously described audio-motor interface and/or top-down prediction processes, not only in neural entrainment to statistical regularities, but also in the formation of explicit knowledge from this implicitly available information.

While our ITC results indicate that neural entrainment happens very fast, the results cannot pinpoint exactly when learning at the neural level is transformed into explicit, consciously accessible knowledge. However, as pointed out in the discussion of our behavioral results, prior findings suggest that the length of exposure may critically influence whether participants are eventually able to explicitly recognize the tone sequences. Altogether, these results point to rapid sensitivity of specific cortical brain regions to hidden statistical structure, which may then be followed by the gradual emergence of knowledge that can be expressed explicitly, occurring in a majority of participants but not all.

#### 4.4. Limitations and Future Directions

The current study used a rather short exposure time, and it is not clear whether a longer exposure time would allow all participants to achieve explicit knowledge of the underlying structure. In addition, we did not control for the attentional level of the participants, so we are not able to determine the influence of attentional control on the process. To test the role of exposure time and attention on statistical learning, future studies that systematically manipulate these variables – like the study by Batterink and Paller (2019), who compared statistical learning under conditions of full and divided attention – will be necessary. Gaining further insights into the dynamic transition from an implicit, neural representation of structure towards an explicitly available one may be used to optimize learning strategies, both in the auditory domain as well as in other modalities. Furthermore, these results could lead to the development of new strategies for individuals with learning impairments.

## 5. Conclusions

MEG is well suited to detect nonlinguistic statistical learning at both the sensor and source level. Combined with behavioral data, these results shed light on different aspects of statistical learning. The present study showed that a wide range of brain areas are involved in this process of learning, not only those primarily related to auditory functions. It also provided evidence that while the learning effects at the neural level are quite robust and strong, the more explicit expression of this kind of nonlinguistic learning is difficult for participants. Nevertheless, these two components are systematically linked, as shown by the correlational results. The nature and timing of these relations suggest that sufficient exposure is necessary to build on the implicit encoding of an underlying structure represented by neural entrainment and to form explicitly available knowledge.

## 6. Acknowledgements

We want to thank Jürgen Dax for his support with the technical setup. This work was partially funded by the FET Open Luminous project (H2020 FETOPEN-2014-2015-RIA under agreement No. 686764) as part of the European Union’s Horizon 2020 research and training program 2014–2018 and the German Federal Ministry of Education and Research (BMBF) to the German Center for Diabetes Research (DZD e.V. 01GI0925).

## 7. Author Contributions

*Julia Moser:* Conceptualization, investigation, methodology, formal analysis and writing – original draft. *Laura Batterink:* Conceptualization, methodology, formal analysis and writing – original draft. *Yiwen Li Hegner:* Methodology and formal analysis. *Franziska Schleger:* Conceptualization, methodology and writing – review & editing. *Christoph Braun:* Methodology and supervision. *Ken Paller:* Conceptualization and writing – review & editing. *Hubert Preissl:* Supervision, conceptualization and writing – review & editing. All authors read and approved the final version of the manuscript.

